# *DE NOVO* SEQUENCING AND ANALYSIS OF THE *RANA CHENSINENSIS* TRANSCRIPTOME TO DISCOVER PUTATIVE GENES ASSOCIATED WITH POLYUNSATURATED FATTY ACIDS

**DOI:** 10.1101/2020.03.10.985457

**Authors:** Jingmeng Sun, Zhuoming Wang, Weiyu Zhang

**Author notes:** **Corresponding author:** Weiyu Zhang, College of Pharmacy, Changchun University of Chinese Medicine, 130117, Changchun, Jilin, China.

## Abstract

*Rana chensinensis* (*R. chensinensis*) is an important wild animal found in China, and a precious animal in Chinese herbal medicine. *R. chensinensis* is rich in polyunsaturated fatty acids (PUFAS). However, information regarding the genes of *R. chensinensis* related to the synthesis of PUFAs is limited. To identify these genes, we performed Illumina sequencing of *R. chensinensis* RNA from the skin and Oviductus Ranae. The Illumina Hiseq 2000 platform was used for sequencing, and the I-Sanger cloud platform was used for transcriptome *de novo* sequencing and information analysis to generate a database. Through the database generated by the transcriptome and the pathway map, we found the pathway for the biosynthesis of *R. chensinensis* PUFAs. The Pearson coefficient method was used to analyze the correlation of gene expression levels between samples, and the similarity of gene expression in different tissues and the characteristics in their respective tissues were found. Twelve differentially expressed genes of PUFA in skin and Oviductus Ranae were screened by gene differential expression analysis. The 12 unigenes expression levels of qRT-PCR were used to verify the results of gene expression levels consistent with transcriptome analysis. Based on the sequencing, key genes involved in biosynthesis of unsaturated fatty acids were isolated, which established a biotechnological platform for further research on *R. chensinensis*.

## INTRODUCTION

*Rana chensinensis* (*R. chensinensis*) is an important wild animal in China. Oviductus Ranae, a valuable Chinese crude drug, is recorded in Pharmacopoeia of the People’s Republic of China as a dried oviduct of the female Chinese frog [9], *R. chensinensis*, distributed mainly in northeastern China. Oviductus Ranae is an established and highly valued food and medicine. Traditional Chinese medicine holds that Oviductus Ranae can moisten the lungs, nourish yin, and replenish the kidney essence [3]. Meanwhile, modern pharmacological studies have demonstrated the activity of Oviductus Ranae in improving immunity, as well as its anti-fatigue, anti-oxidative, anti-lipemic, and anti-aging properties [10]. Oviductus Ranae has an established safety profile, it is a raw material with natural health care functions, and has great potential for further use, therefore, it is widely used in food, pharmaceutical and chemical industries. At present, the food developed using Oviductus Ranae involves canned food, candy, yogurt and beverages. Moreover, there are various administration forms (i.e., pills, capsules, and granules) produced from Oviductus Ranae. In the skin care industry, the active ingredients (i.e., unsaturated fatty acids, carotene, and vitamins) in Oviductus Ranae can help improve skin dryness, reduce pigmentation, and offer a cosmetic effect [11].

*R. chensinensis* is a cold-tolerant vertebrate amphibian that grows for ≤6 months in hibernation [12]. Maintaining the fluidity of the cell membrane in a low-temperature environment ensures that it can perform its normal physiological functions [8]. It is known to all that the fluidity of the cell membrane is closely related to the composition of polyunsaturated fatty acids (PUFAs), the content of PUFAs in the cell membrane is very important for maintaining cell structure, membrane mobility, and enzymatic activity. PUFAs cannot be ingested from the external environment by hibernating animals. Therefore, we investigated the mechanism involved in the survival of *R. chensinensis* during hibernation and changes in the content of PUFAs. We believe that PUFAs, which are abundantly in *R. chensinensis*, may be the reason for the decrease in fatty acid saturation by *R. chensinensis* in the low-temperature environment. The synthetic pathway is the presence of fatty acid desaturase (FADS) in the organism, which is a key enzyme in the synthesis of PUFAs.

There are four main kinds of FADS in animals, namely Δ9-FAD, Δ5-FAD, Δ6-FAD, and Δ4-FAD [8]. Of those, Δ6-FAD and Δ5-FAD are the first and second rate-limiting enzymes. Studies have found that a low-temperature environment can cause up-regulation of Δ9-FAD gene expression. Previous experimental studies have found significant differences in fatty acid content in Oviductus Ranae collected in different seasons. The content of PUFAs in the predation growth period and scattered hibernation samples was 14.16% and 29.83%, respectively. Therefore, we hypothesized that FADs is necessary for the synthesis of PUFAs in *R. chensinensis*, which affect their own synthesis of PUFAs under low-temperature stimulation. At present, genetic information regarding *R. chensinensis* remains unknown, and the molecular mechanism of fatty acid synthesis in *R. chensinensis* is unclear [7]. Therefore, we used non-reference transcriptome sequencing technology to obtain the genetic information of *R. chensinensis*. The FADs gene *in vivo* was identified by studying the changes in the content of PUFAs in *R. chensinensis*. Through the detection of FADs gene expression in Oviductus Ranae and the skin of *R. chensinensis*, the role of this gene in the synthesis of PUFAs was elucidated, and the pathway of PUFA synthesis was determined.

## MATERIALS AND METHODS

### Animals and treatments

To ensure the space-time specificity of the sample, We removed Oviductus Ranae and skin from R. chensinensis, rapidly frozen in liquid nitrogen, and stored in an ultra-low temperature freezer at −80°C. All procedures performed in this study involving the handling of *R. chensinensis* were approved by the Animal Care and Welfare Committee of Changchun University of Chinese Medicine (Jilin, China).

### RNA isolation and reverse transcription complementary DNA (cDNA)

RNA was extracted from the skin and Oviductus Ranae of *R. chensinensis*. Detection of RNA concentration and quality was performed using Nanodrop2000 (Thermo Scientific, U.S.A.). Total RNA integrity was determined through 1.2% agarose gel electrophoresis. Sample reverse transcription was performed using Takara’s (Takara, China) PrimeScript™ RT reagent Kit with gDNA Eraser-Perfect Real Time Kit (Code No. RR047A). The reaction system include the following: reaction solution 10.0μL, 5×PrimeScript buffer 10.0μL, PrimeScript™ RT Enzyme Mix I 1.0μL, RT primer mix 1.0 μL, and Rnase free dH_2_O 4.0μL, in a total volume of 20μL. The reaction procedure was: 37°C for 15min, followed by 85°C for 5s. The obtained cDNA was stored at −20°C. Transcriptome sequencing was performed using the Illumina Hiseq 2000. The data were analyzed on the free online platform of Majorbio I-Sanger Cloud Platform (www.i-sanger.com). De novo transcriptome assembly was carried out using the Trinity software (https://github.com/trinityrnaseq/trinityrnaseq) [1].

### De novo assembly and comparative analysis between two samples

Using the Trinity software to head assembly of all the clean data, we spliced the transcript sequence (i.e., the longest transcript of each gene, defined as unigene), as a basis for the follow-up bioinformatics analysis. The TransRate (http://hibberdlab.com/transrate/) software of the transcriptome assembly sequence filter was used and optimized from the beginning. The CD-HIT (http://weizhongli-lab.org/cd-hit/) software and the sequence alignment Cluster method were used to remove redundancy and similar sequences, and finally obtain the non-redundant (NR) sequence. BUSCO (Benchmarking Universal single-copy Orthologs, http://busco.ezlab.org) evaluates the assembly integrity of the genome or transcriptome using single copy straight homologous genes. Genome assembly required TBLASTN comparison with the consistent sequence of BUSCO. Subsequently, Augustus was used to predict the genetic structure, and finally, HMMER3 comparison was used.

### Identification of differentially expressed genes (DEGs)

The fragments per kilobase million (FPKM) algorithm was used to quantify the abundance of the transcript in the DEG analyses [6].The DEGs were identified using the DESeq2 (http://bioconductor.org/packages/stats/bioc/DESeq2/),DEGseq(http://bioconductor.org/packages/stats/bioc/DEGSeq/), and edgeR [13]. For experimental designs with biological replicates, the raw counts were statistically analyzed directly using the DESeq2 software based on the negative binomial distribution. Genes for comparing differences in expression between groups were obtained based on certain screening conditions. The default parameter was p-adjusted to <0.05 and |log2FC| ≥1. A p-value ≤0.001 and a>2-fold change (absolute value of log_2_ ratio>1) in gene expression denoted statistical significance.

### Functional annotation and analysis of pathway enrichment

The assembled transcriptome sequences were compared with those in the NR (ftp://ftp.ncbi.nlm.nih.gov/blast/db/),Swiss-Prot(http://web.expasy.org/docs/swiss-prot_guideline.html), Pfam (http://pfam.xfam.org/) [2], Clusters of Orthologous Groups (COG of proteins; http://www.ncbi.nlm.nih.gov/COG/), Gene Ontology (GO; http://www.geneontology.org), and Kyoto Encyclopedia of Genes and Genomes (KEGG; http://www.genome.jp/kegg/) databases to obtain the annotation information for each database. Subsequently, the annotation information for each database was calculated. By comparing with the KEGG database, the KO number corresponding to the gene or transcript was obtained, According to the KO number, the specific biological pathway involved in the gene or transcript can be determined. Functional annotation, categorization, and protein evolution analysis can be performed by comparison with the COG database. By comparing with the NR library, the similarity of the transcript sequence of the species to other species and the functional information of the homologous sequence can be obtained.

### Real-time fluorogenic quantitative PCR

This experiment used Takara’s SYBR Premix EX Taq™ (Tli RNaseH Plus) kit, and its quantitative part was performed on the Mx3000P™ (Agilent Technologies, CA, U.S.A.) Real time PCR instrument. Its operating system is Stratagene (Mx3000P). Three replicates and negative controls were set for each sample. In this experiment, GeNorm software was used to screen the housekeeping genes in the samples, and the stability of five candidate internal reference genes EF1α, GAPDH, EPB2, TUB and ACT were analyzed. The results showed that the internal reference gene EF1-α had the highest comprehensive stability, so EF1α was selected as the internal reference gene of the experiment. The most stable housekeeping reference gene EF1-α was selected for the expression analysis in various tissues. In the obtained gene database, we screened 12 PUFAs genes by differential gene expression. Their expression levels are significantly different in the skin and Oviductus Ranae to verify that the gene expression levels of the transcriptome analysis are consistent. The relative expression of twelve genes was normalized to the expression of EF1α and expressed relative to the level in various treatment. Primer express design software was used to design primers based on Blast analysis of 12 differential genes and specific regions of internal reference genes (Primer sequence see attachment). The optimized reaction system included the following: TransStart Top Green qPCR Super Mix (2×) 10μL, Passive Reference Dye (50×) 0.4μL, PCR forward primer (10μm) 0.4μL; PCR reverse primer (10μm) 0.4μL; old H_2_O 6.8μL; and cDNA 2μL, in a total volume 20μL. The two-step PCR amplification standard procedure was as follows: pre-denaturation 95°C, 30s; PCR reaction 95°C, 5s; 60°C, 15s; 40 cycles; dissolution curve 95°C, 5s; 60°C, 60s; 95°C, 15s. The fold change in relative expression level was calculated using 2^−△△CT^ method [5].

## RESULTS

### Transcriptome sequencing and de novo assembly

In this study, RNA-seq technology was used to investigate the transcriptome in Oviductus Ranae and skin samples obtained from *R. chensinensis*. Six cDNA libraries were constructed, representing Oviductus Ranae and skin, respectively. More than 93% of the data yielded a high-quality score. In total, 338843554nt bases were generated. The results of the assembly yielded 305,087 unigenes; the average length was 608.81nt and the N50 was 865nt.

### Functional annotation of unigenes

The assembled transcriptome sequences were compared with those in six databases (NR, Swiss-Prot, Pfam, COG, GO, and KEGG) to obtain annotation information for each database. Statistical analyses on the annotations for each database were performed. The BLAST search revealed that 26.29% of the unigenes exhibited a significant match to genes in the NR database, followed by 23.11% in the Swiss-Prot, 18.92% in the Pfam,16.69% in the KEGG, 10.84% in the GO, and 7.87% in the COG databases (Table 1).

**Table 1.**
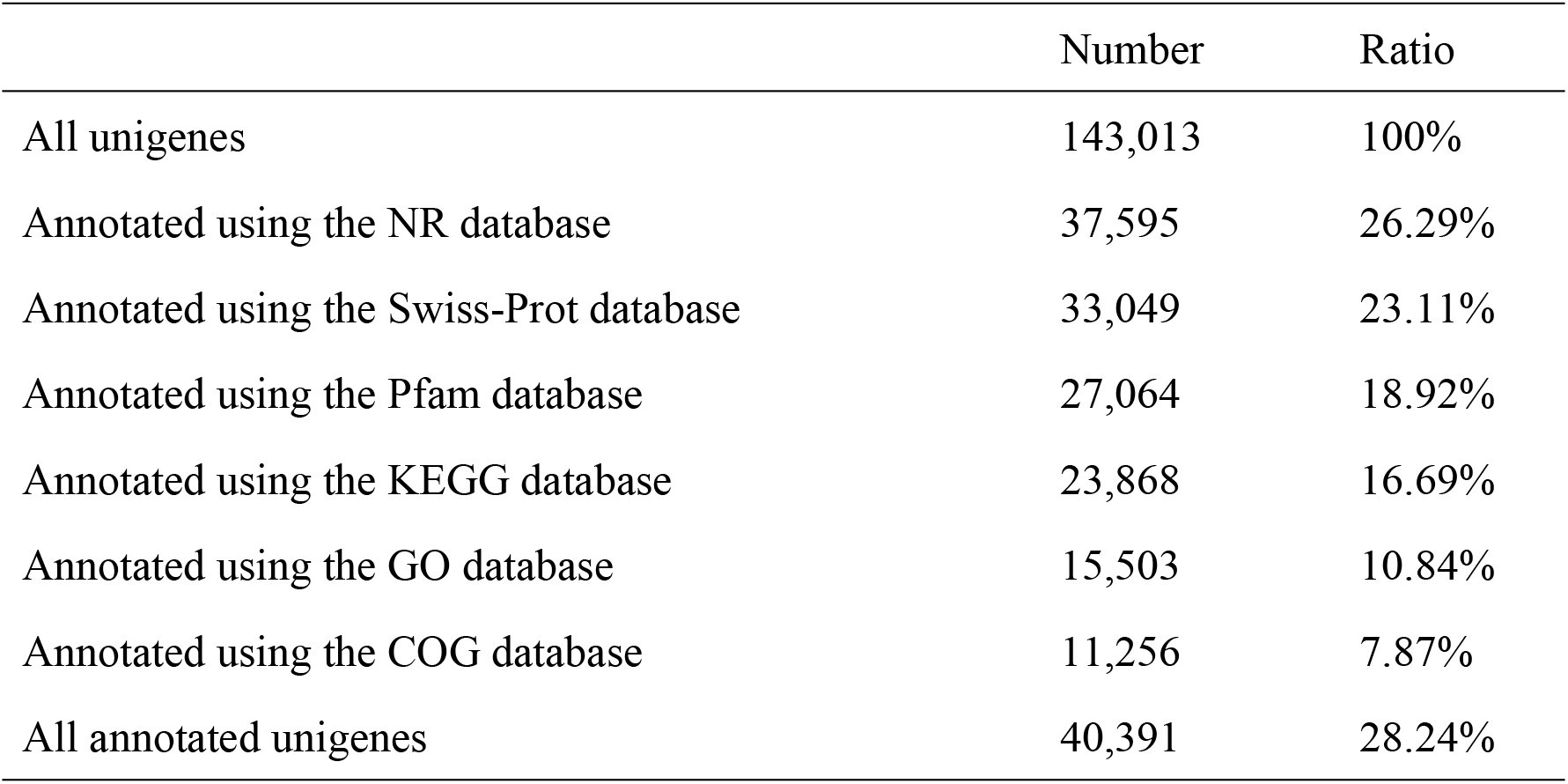
Summary of all unigenes annotated in the Oviductus Ranae and skin

**Table 2:** The NCBI number corresponding to the 12 selected genes

| Unigene | NCBI |
| --- | --- |
| unigene97926 | XP 012819853.1 |
| unigene97708 | ACO51759.1 |
| unigene113904 | NP 001107300.1 |
| unigene112919 | XP 002932858.1 |
| unigene99521 | NP 001091202.1 |
| unigene104803 | XP 012819853.1 |
| unigene100548 | XP 002934815.1 |
| unigene97226 | NP 001086822.1 |
| unigene90014 | AAH73571.1 |
| unigene106327 | XP 012808582.1 |
| unigene105430 | XP 010383225.1 |
| unigene111094 | XP 002940372.2 |

**Table 3:** 12 genes physical and chemical properties

| Unigene | AA | MW | pI | Aliphatic index | GRAVY |
| --- | --- | --- | --- | --- | --- |
| 97708 | 459 | 53728.89 | 8.91 | 85.16 | -0.167 |
| 111094 | 207 | 23635.12 | 8.91 | 112.56 | 0.419 |
| 105430 | 192 | 19954.36 | 10.52 | 39.79 | -0.692 |
| 106327 | 522 | 59634.71 | 9.68 | 76.78 | -0.285 |
| 90014 | 255 | 28318.19 | 9.65 | 84.51 | -0.065 |
| 99521 | 698 | 79329.22 | 8.23 | 66.2 | -0.375 |
| 97226 | 587 | 68623.3 | 10.06 | 93.99 | -0.011 |
| 112919 | 749 | 81919.3 | 9.58 | 99.33 | 0.334 |
| 113904 | 752 | 84364.37 | 8.81 | 82.21 | -0.068 |
| 104803 | 505 | 59278.41 | 9.25 | 95.49 | 0.119 |
| 100548 | 635 | 75126.35 | 9.42 | 81.65 | -0.3 |
| 97926 | 252 | 29585.1 | 9.62 | 90.95 | 0.136 |

AA means number of amino acids; MW means molecular weight; pI means theoretical pI; GRAVY means grand average of hydropathicity.

The threshold E-value of the annotated unigenes against the NR database was 1e-5. Only 16.2% of the unigenes exhibited strong similarity (<1e-100) with the sequence in the NR database, whereas the E-values for 83.9% of the unigenes ranged from 1e-5 to 1e-100 (Figure 1A). The distribution of similarity was as follows: >80%, 60–80%, and 40–60% for 34.3%, 28.7%, and 22.4% of the sequences, respectively(Figure 1B). For species distribution matched against the NR database, 49% of the matched unigenes showed similarities with *Silurana tropicalis*, followed by the African clawed frog (15.4%) (Figure 1C).

**Figure 1.**
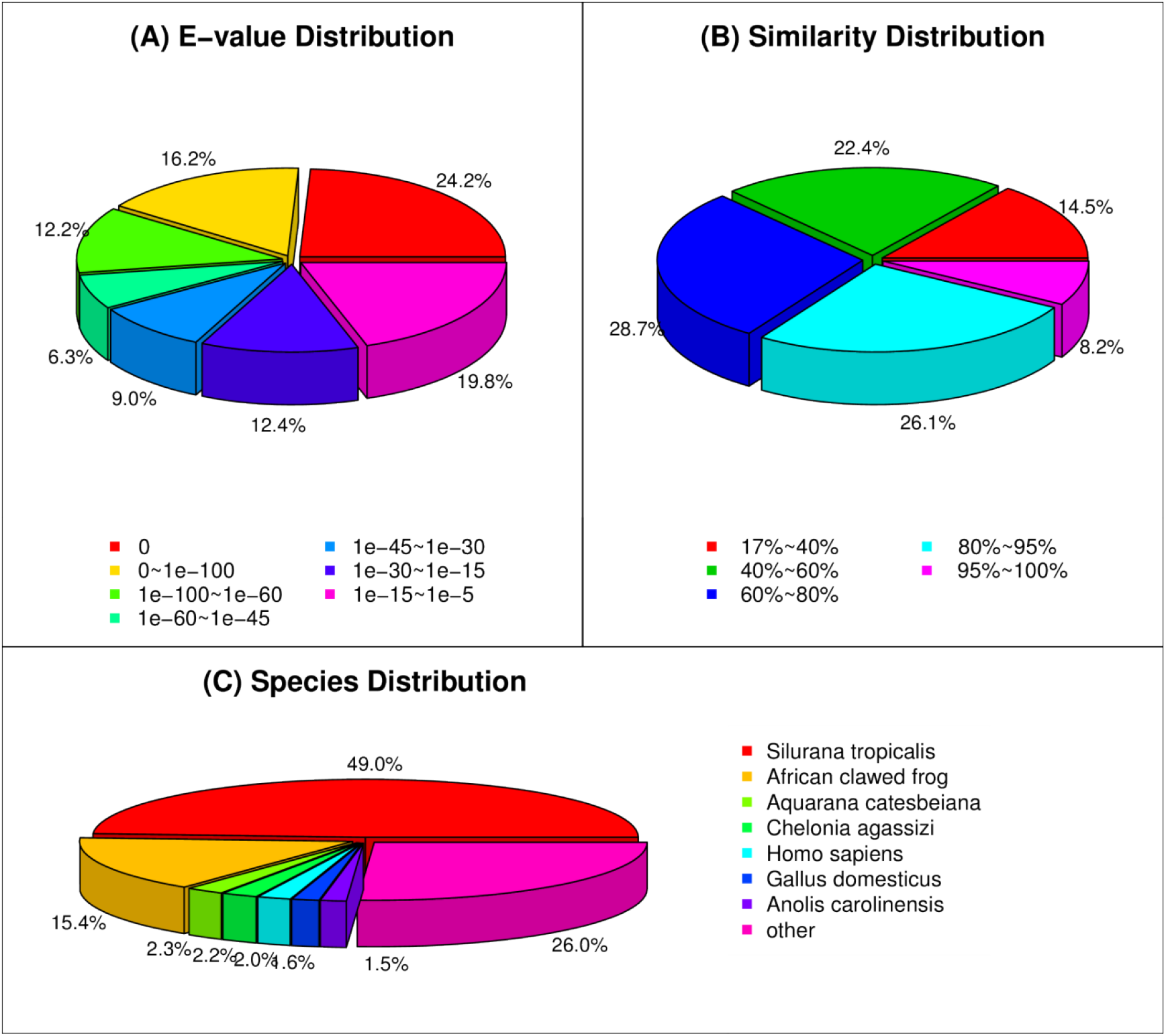
NR classification

The Unigene and COG databases were compared to predict the possible functions of unigenes and perform functional classification statistics (Figure 2). The hits from the COG prediction were functionally classified into 25 categories, in which the most enriched terms were general function prediction only (8,073 unigenes, 20%), followed by replication, recombination and repair (3,538 unigenes, 9%), and transcription (2,934 unigenes, 7%). It is indicated that 20% of unigenes in *R. chensinensis’s* skin and Oviductus Ranae function as general function prediction only. The least unigenes function is extracellular structures and nuclear structure, but it does not mean that they can not play this role.

**Figure 2.**
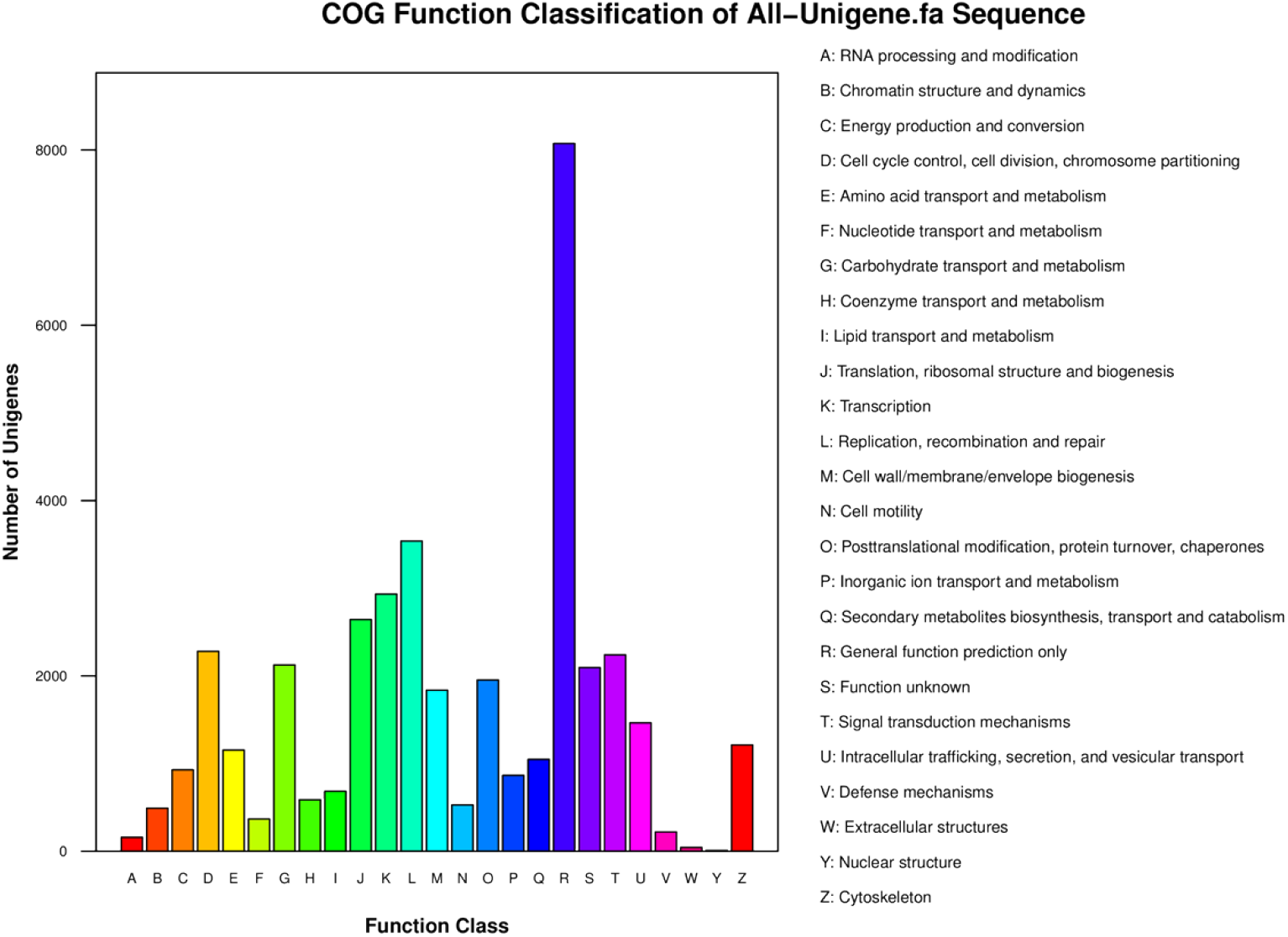
COG function classification of unigenes in All-Unigene

GO analysis regarding the putative proteins was performed using Blast2GO. GO is an internationally standardized gene functional classification system, which comprehensively describes the properties of genes and gene products in organisms. Unigenes that successfully annotate are classified according to the three independent ontologies of GO, biological processes, cellular components, and molecular functions involved in the gene. Subsequently, functional classification statistics are performed for all unigenes that are annotated in the GO database. The three main GO categories were classified into 56 subcategories. The greatest numbers of transcripts were assigned to biological processes (211,193), cellular components (143,518), and molecular functions (46,271) (Figure 3). Among the biological processes, the greatest number of transcripts was assigned to cellular process (25,480). In cellular components, cells were dominant (25,086). Among the molecular functions, the greatest number of transcripts was assigned to binding (22,978). The distribution of the GO terms showed that cellular process, metabolic process, and single-organism process accounted for the largest proportion of biological processes Moreover, it showed that the cell and cell part were significantly enriched terms among cellular components, and that the binding and catalytic activities were the most represented terms in molecular functions.

**Figure 3.**
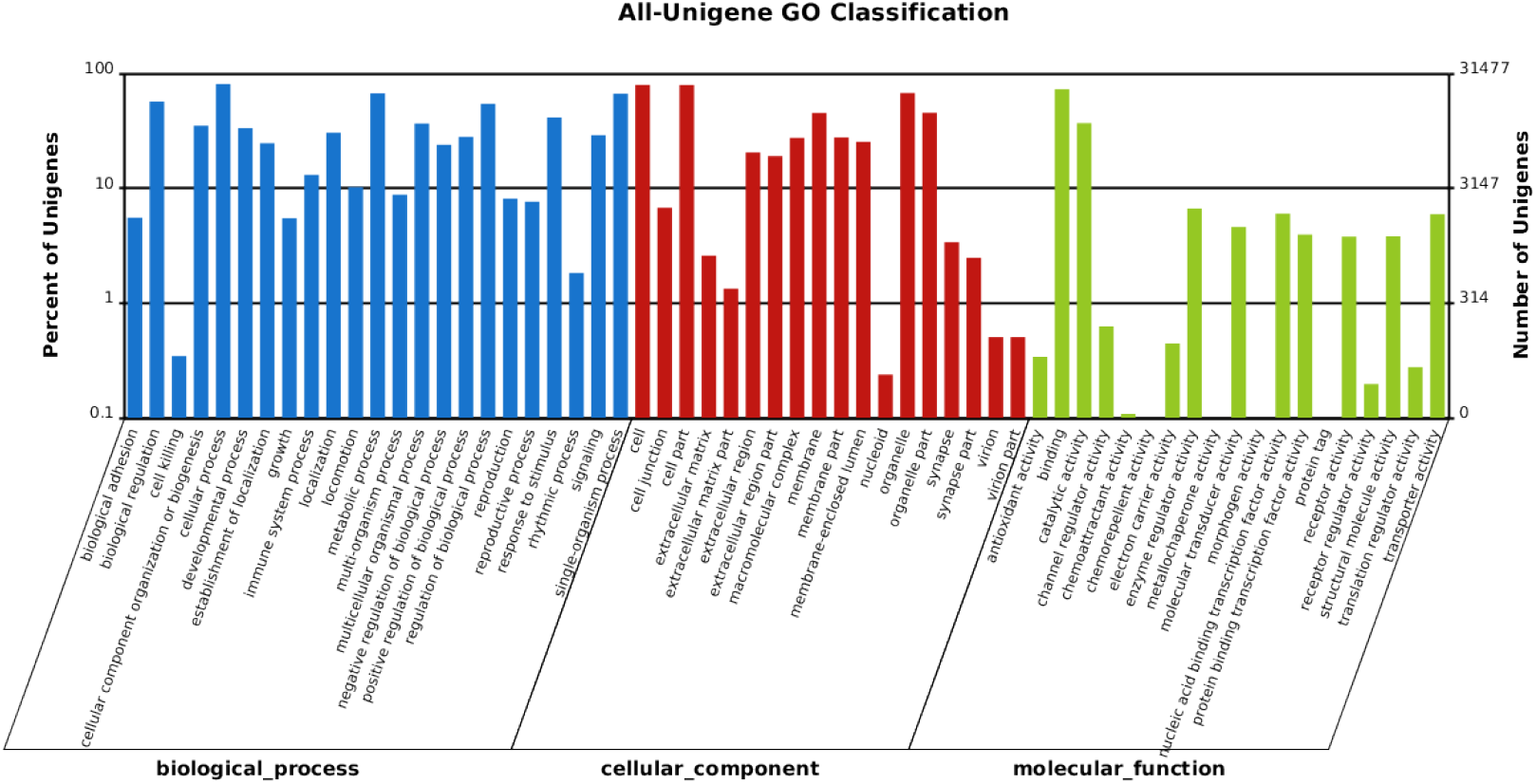
GO classification analysis of unigenes in All-Unigene

Mapping all annotated unigenes to the reference pathway in the KEGG database. In total, 10,569 unigenes were assigned to six clusters and 44 KEGG pathways (Figure 4), including metabolism, genetic information processing, environmental information processing, cellular processes, organismal systems, human diseases. According to the Figure 4, the path with the most unigenes in the environmental information processing is signal transduction. Note that signal transduction is the most important KEGG pathway in environmental information processing. The most popular KEGG pathway category for Unigenes is human diseases. It is shown that human diseases are the most relevant KEGG pathways to *R. chensinensis’s* skin and Oviductus Ranae.

**Figure 4:**
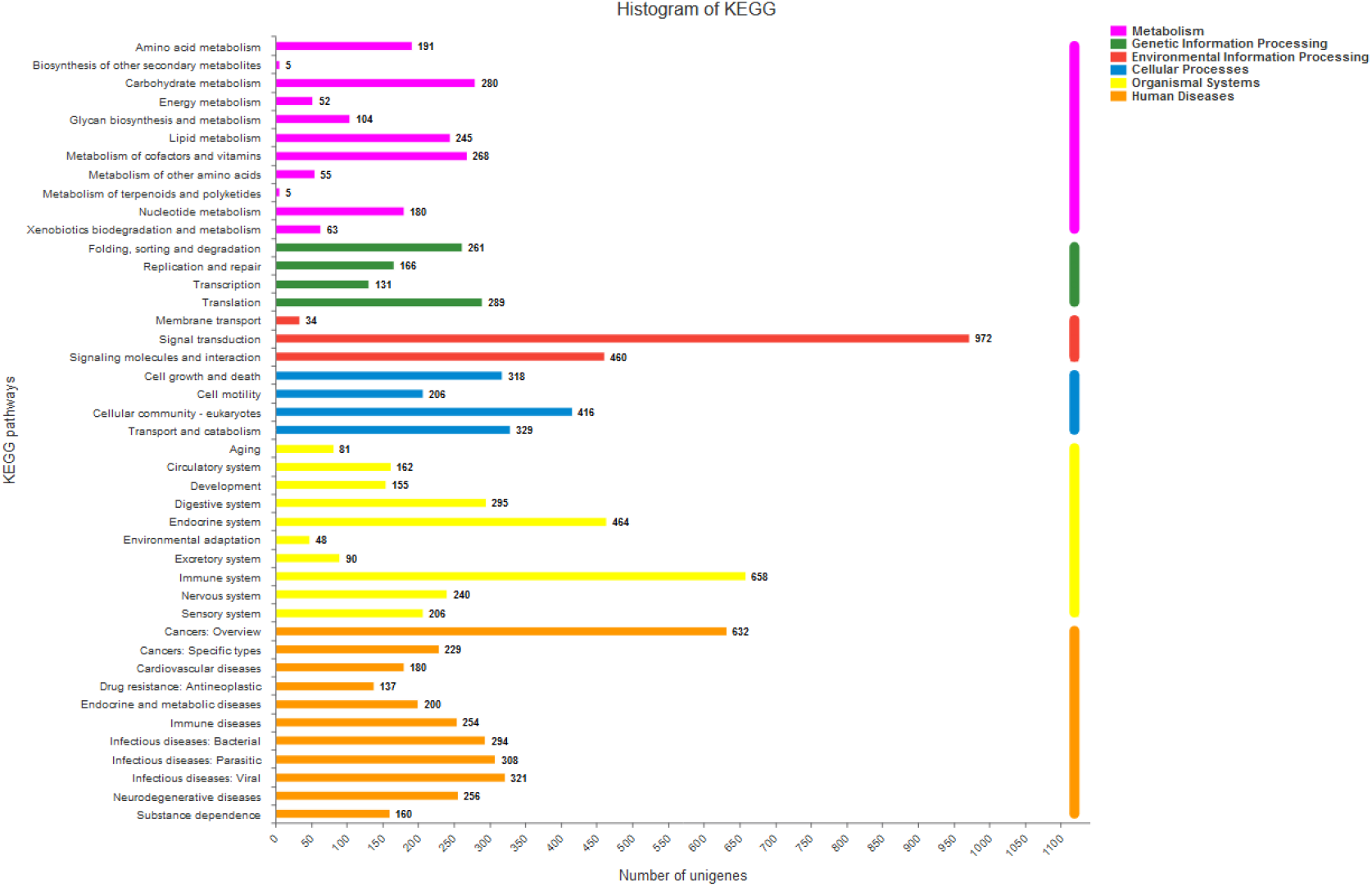
Histogram of the KEGG pathways of assembled unigenes in Oviductus Ranae and skin obtained from *R. chensinensis.* The ordinate is the name of the KEGG metabolic pathway and the abscissa is the number of genes annotated to the pathway. The KEGG metabolic pathway can be divided into 6 categories: metabolism, genetic information processing, environmental information processing, cellular processes, organismal systems, human diseases.

### Differential expression analysis

The FPKM density distribution as a whole reflects the gene expression pattern of each sample. Based on this information, we can check the distribution of unigene FPKM in different tissues of *R. chensinensis* on the whole level, and effectively evaluate the expression of unigenes.

The correlation of gene expression levels between samples is an important index to test the reliability of the experiment and the reasonableness of sample selection. If there is biological duplication in the sample, the correlation coefficient between biological duplication is usually required to be higher. The correlation between samples reflects the degree of similarity between samples, that is, the similarity of the expression levels of samples with different treatments or tissues. A correlation coefficient value close to 1 indicates high similarity and small differences in genes between samples. The correlation coefficient of samples between biological repeats should be greater than that of samples non-biological repeats. There were two groups of samples investigated in this study, and triplicates were set for each group of samples. The Pearson correlation coefficient between samples was calculated using the DESeq2 language. The up-regulated and down-regulated genes identified between the skin and Oviductus Ranae samples were selected. A total of 15, 915 genes showed significant differential expression between the two samples: 7,035 genes were up-regulated, and 8,880 genes were down-regulated. Figure 5 shows the volcano plot for the differential expression level of genes between two samples.

**Figure 5:**
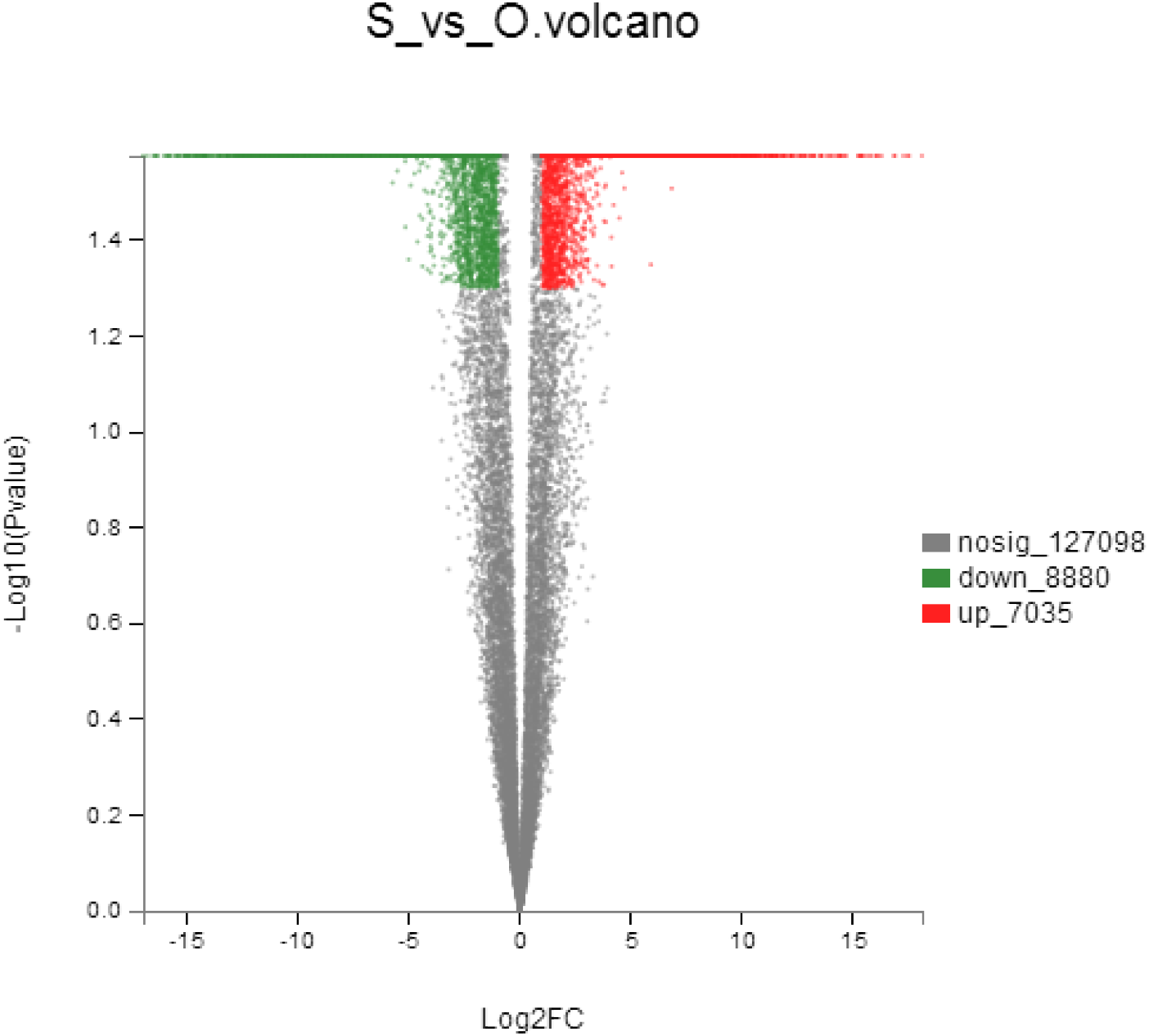
Volcano plot of DEGs in samples of Oviductus Ranae and skin obtained from *R. chensinensis.* S stands for skin. O stands for Oviductus Ranae. The abscissa is the fold change value of the difference in expression of the gene between the two samples, that is, the value obtained by dividing the expression level of the treatment sample by the expression amount of the control sample. The ordinate is a statistical test value for the difference in the change in gene expression, that is, the p value. The higher the p value, the more significant the difference in expression, and the values of the horizontal and vertical coordinates are logarithmically processed. Each point in the figure represents a specific gene. Red dots indicate significantly up-regulated genes, green dots indicate significantly down-regulated genes, and black dots are non-significantly differential genes. After mapping all the genes, it can be known that the point on the left is the gene whose expression is down-regulated, and the point on the right is the gene whose expression is up-regulated. The more the left and upper points are expressed, the more significant the difference.

According to the Venn diagram (Figure 6), it can be seen that the gene expressed in Oviductus Ranae has a total of 25173 unigenes, which represents an immune function. The gene expressed in the skin a total of 34421 unigenes, representing antioxidant function. The skin and Oviductus Ranae coincide with a total of 29023 unigenes, accounting for about 25% of the total, indicating that the coincident part has both immune function and antioxidant activity.

**Figure 6:**
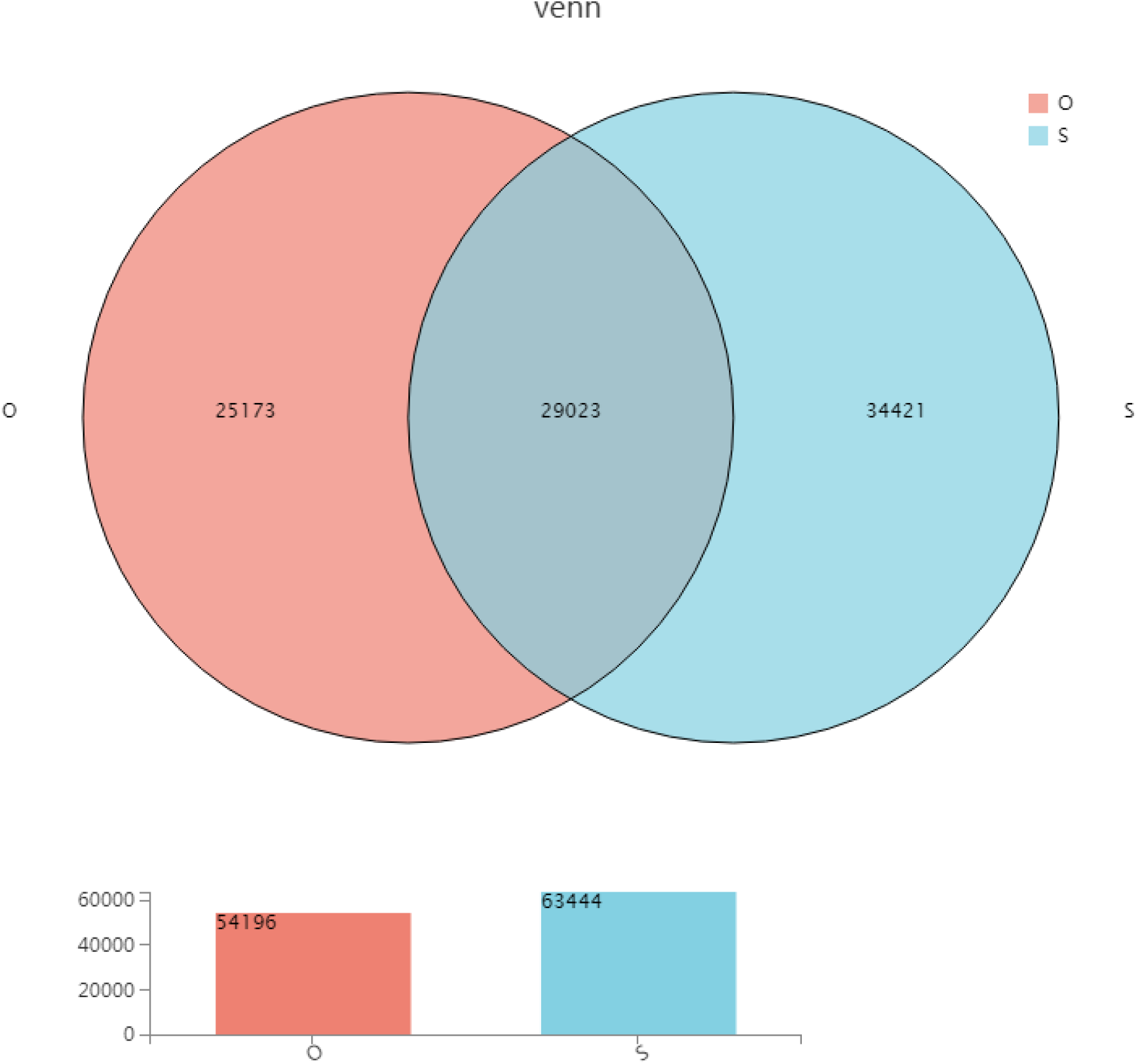
Venn diagram of DEGs in samples of Oviductus Ranae and skin obtained from *R. chensinensis.* O stands for Oviductus Ranae, S stands for skin. Venn diagram between samples: circles of different colors represent the number of unigenes expressed in a set of samples. The intersecting area of the circle represents the number of unigenes shared by each group. Column chart: the abscissa indicates the sample name and the ordinate indicates the number of expression unigene.

Differential expression analysis was performed to identify genes with different expression levels between different samples. Moreover, GO function analysis and KEGG pathway analysis on differentially expressed genes were conducted. In Oviductus Ranae and skin obtained from *R. chensinensis*, functional categories were linked to various metabolisms and biosynthesis. In Figure 7, pathways involved in glycosphingolipid biosynthesis - lacto and neolacto series, tyrosine metabolism, linoleic acid metabolism, drug metabolism - cytochrome P450, arachidonic acid metabolism, hematopoietic cell lineage, and pancreatic secretion were enriched.

**Figure 7:**
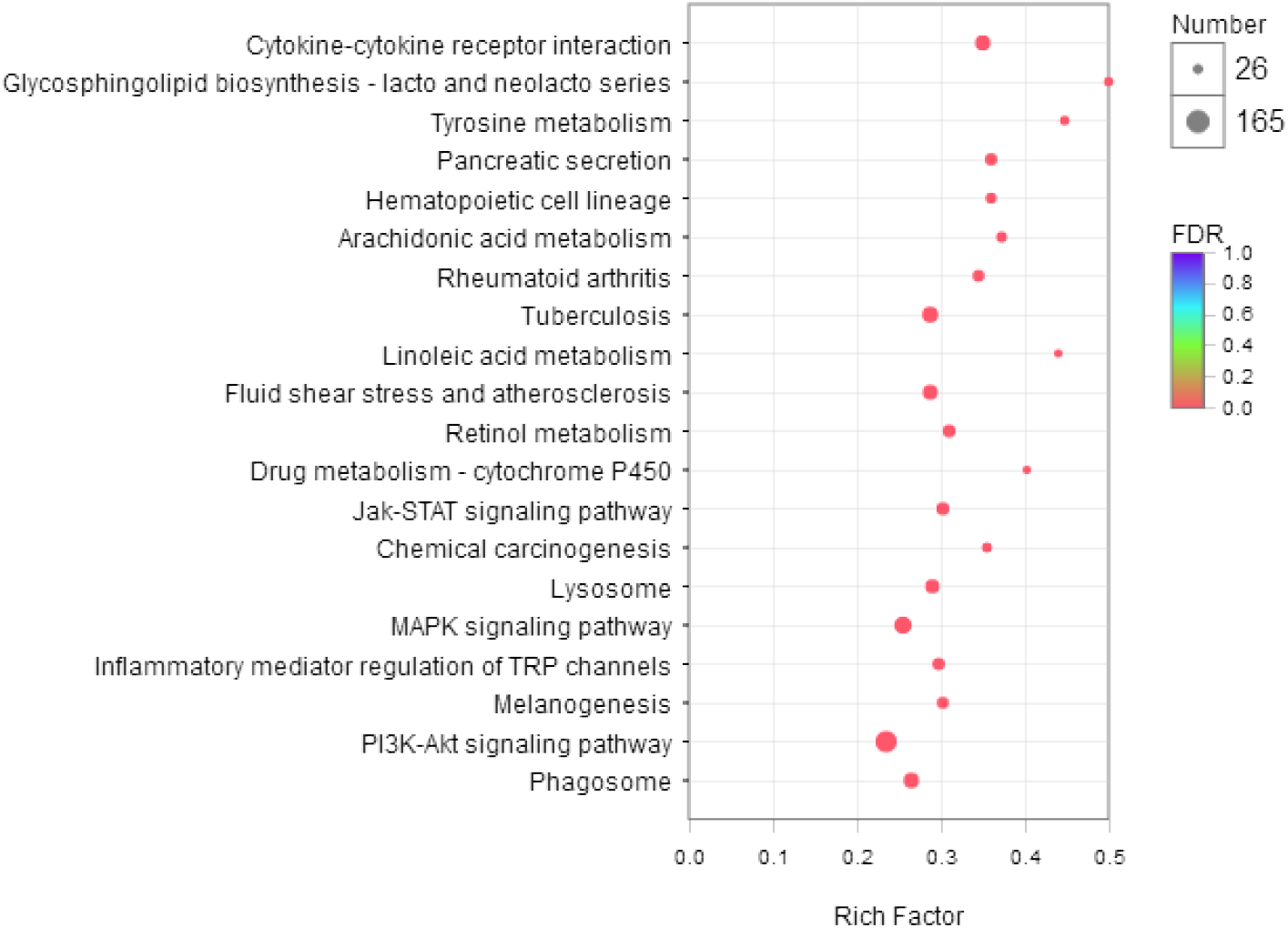
Scatterplot of the KEGG pathway enrichment analysis of differential expressed genes in paired comparisons of Oviductus Ranae and skin obtained from *R. chensinensis.* The vertical axis represents the path name and the horizontal axis represents the Rich factor [The ratio of the unigene number (Sample number) enriched in the path to the annotation unigene number (Background number). The larger the Rich factor, the greater the degree of enrichment.] The size of the point indicates how many genes are in the path, and the color of the point corresponds to a different Qvalue range.

### Phylogenetic analysis of key genes in the biosynthesis of fatty acids

Multiple alignments of the full-length bZIP sequences of the Rana chensinensis gene were performed using the ClustalX 2.0 program, with the parameters set to default and saved in the ClustalX file format. The comparison file is input into the MEGA 7.0 program to build a phylogenetic tree. The construction method is Neighbor-Joining. The specific parameters are set to: p-distance model, and the bootstrap value is 1000. Genes annotated as *R. chensinensis* fatty acids in the transcriptome were shortlisted. The obtained 12 sequences of *R. chensinensis* PUFAs related genes were translated into amino acid sequences. BLAST alignment was performed in the National Center for Biotechnology Information (NCBI) platform, and the results of DNA-man comprehensive alignment were analyzed. The data showed that *R. chensinensis* exhibited the highest homology with *Genus Nanorana*, *Rana catesbeiana*, and the African clawed frog. The sequences of these three species were downloaded from the NCBI platform, and the MEGA 5.0 was used to construct the phylogenetic tree (Figure 8).

**Figure 8:**
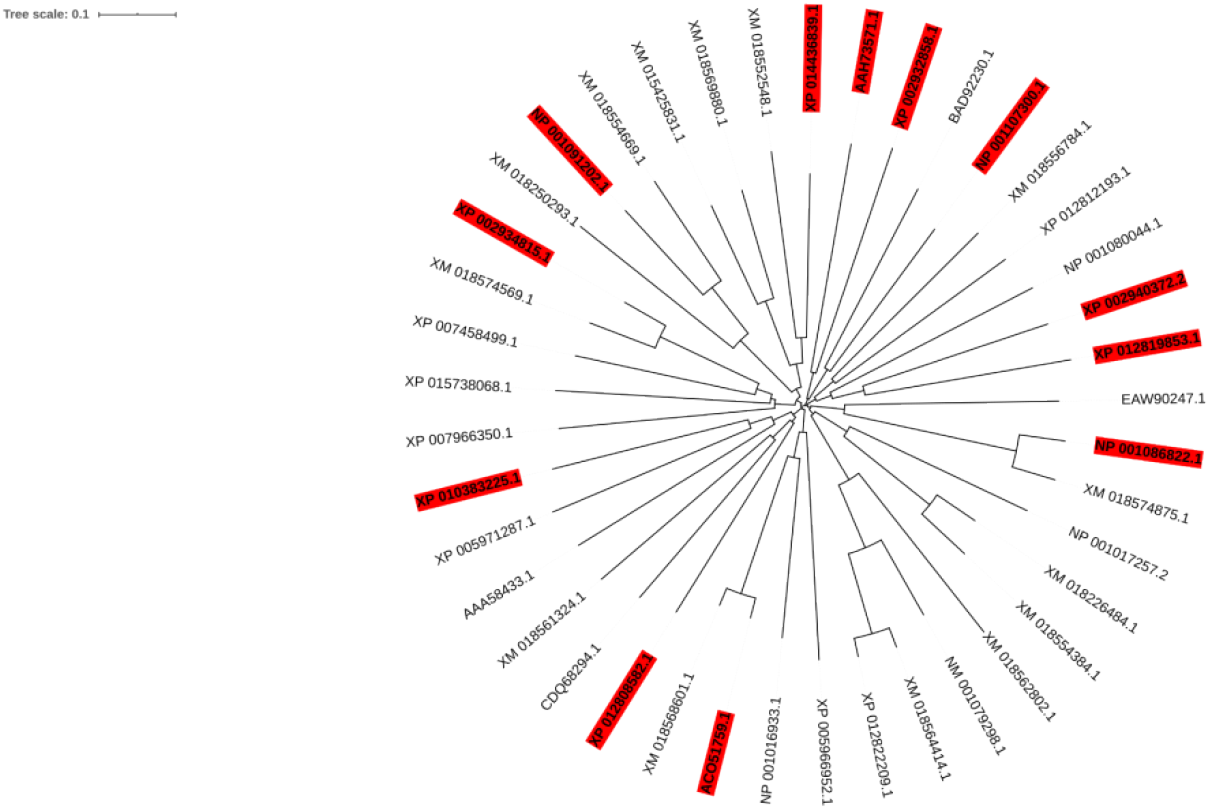
The phylogenetic tree was constructed with the MEGA 5.0 software using the neighbor-joining method (in the red part, 12 differentially expressed genes were screened for the qRT-PCR validation gene)

### Quantitative real time-PCR (qRT-PCR)

Twelve PUFA unigenes were selected for qRT-PCR assays to confirm the results of the sequencing analysis. The selected unigenes showed differential expression patterns. The results of this investigation were consistent with those observed in the sequencing analysis (Figure 9).

**Figure 9:**
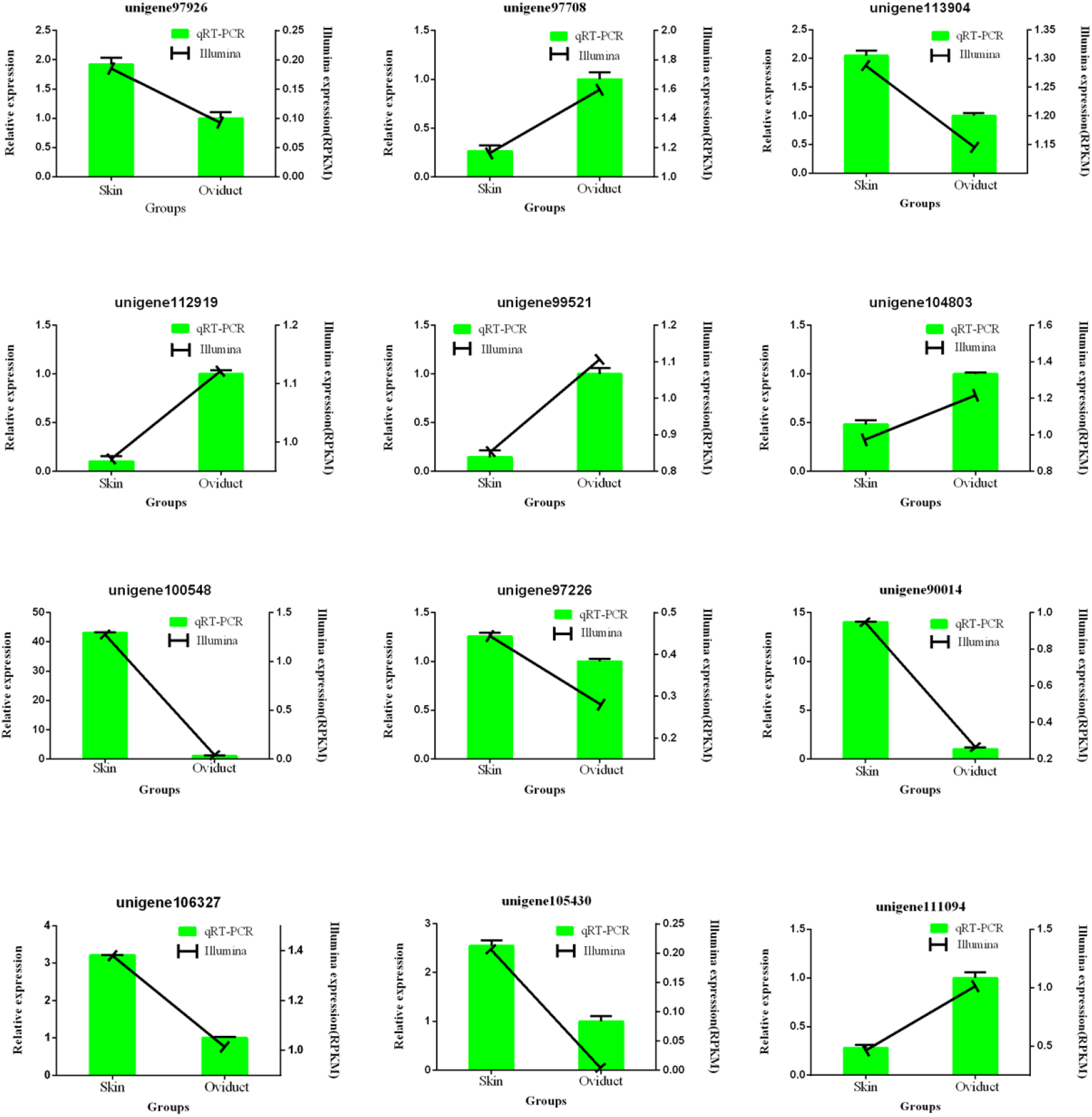
qRT-PCR of the selected unigenes

### Analyses of the unsaturated fatty acids pathway and putative genes in the transcriptome

We focused our analyses on the KEGG pathways and transcripts that appeared to be regulated in the samples to identify unsaturated fatty acids genes (Figure 10). In the biosynthesis of unsaturated fatty acids, we were interested in the two key genes encoding unsaturated fatty acids biosynthetic enzymes, namely long-chain fatty-acyl-CoA hydrolase (EC 3.1.2.2) and Oleoyl-[acyl-carrier-protein] hydrolase (EC 3.1.2.14). The 21 unigenes related to these genes, including those that were up-regulated and down-regulated, are listed in Table 4. Of these, 15 genes were annotated as long-chain fatty-acyl-CoA hydrolase (five up-regulated and 10 down-regulated), while six were annotated as oleoyl-[acyl-carrier-protein] hydrolase (four up-regulated and two down-regulated).

**Table 4:**
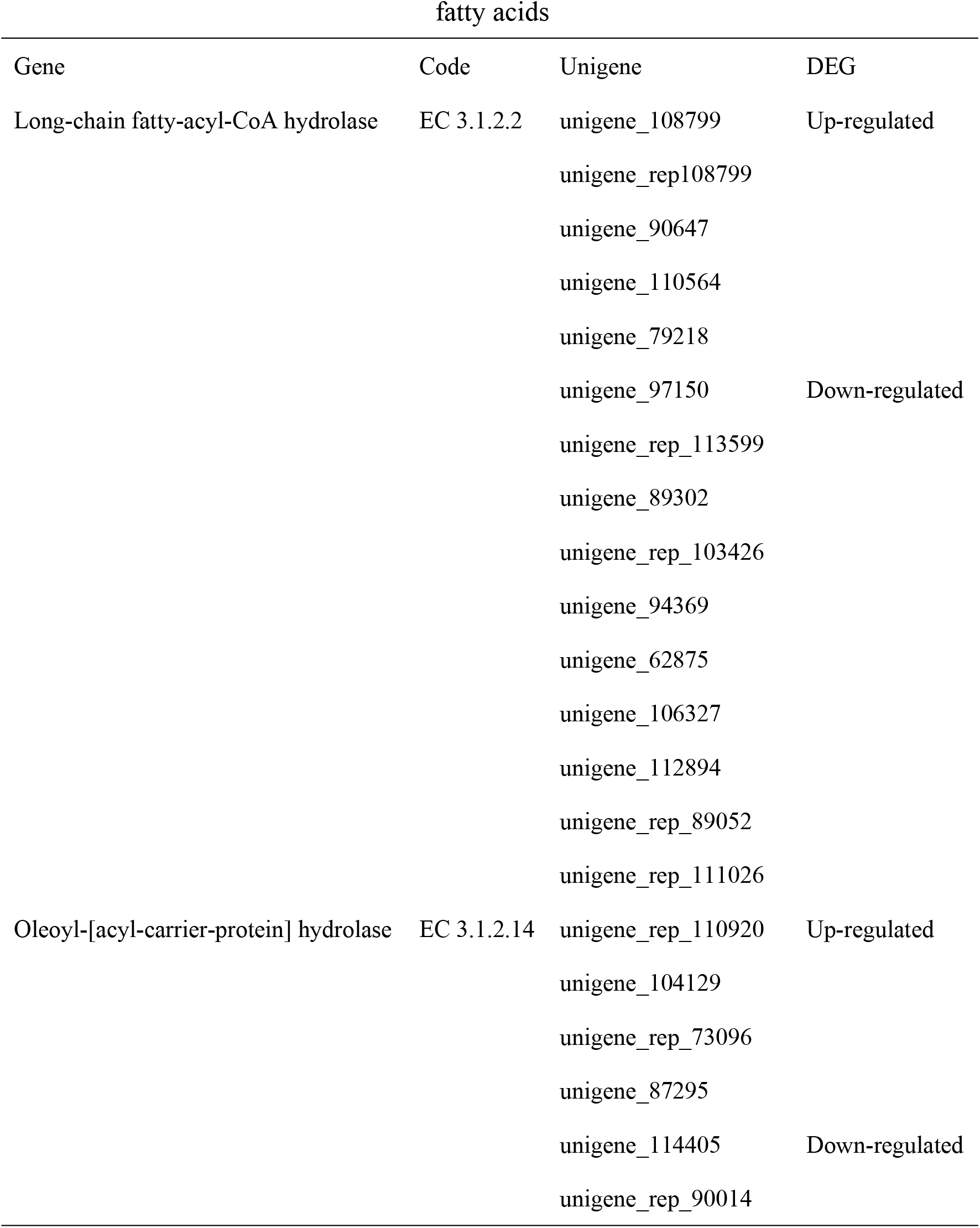
Unigenes predicted to be associated with the biosynthesis of unsaturated fatty acids

**Figure 10:**
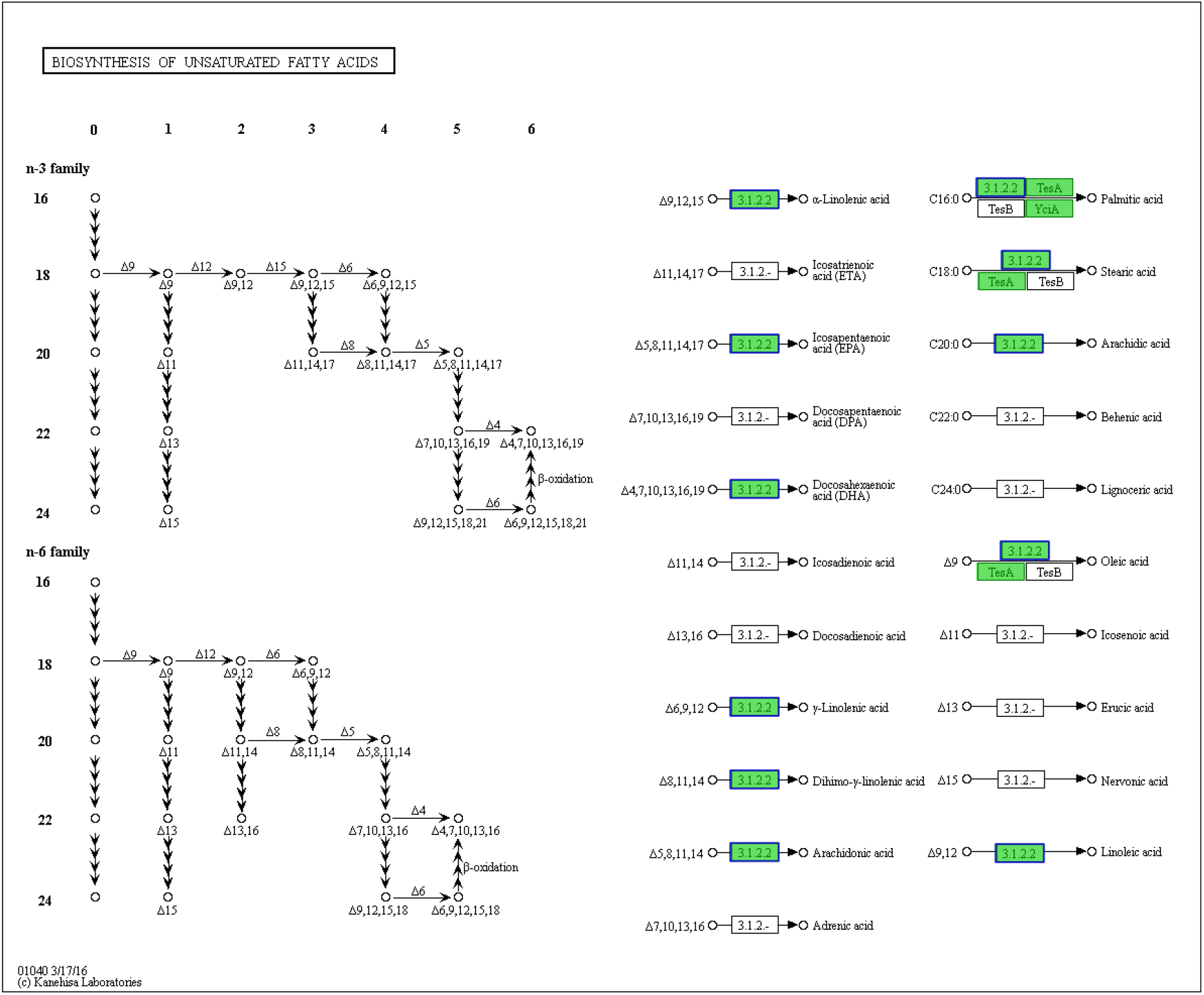
The metabolic pathways for the biosynthesis of unsaturated fatty acids

## DISCUSSION

*R. chensinensis* has been applied in Chinese herbology to resist sickness and enhance immunity, owing to its anti-inflammatory, anti-fatigue, and antioxidant properties [14]. In Northeast China, the artificial feeding quantity of *R. chensinensis* increases annually. According to incomplete statistics, in 2018>600 million *R. chensinensis* frogs were harvested in the Jilin province (one of the provinces in the northeast of China) [15]. In the process of Oviductus Ranae synthesis, it is mainly the accumulation of fatty acids. During the accumulation of fatty acids, genes of key enzymes determine the synthesis of unsaturated fatty acids. Therefore, we need to study the changes and accumulation of fatty acids at the genetic level. However, owing to the lack of genetic resources of *R. chensinensis*, we adopted a transcriptome sequencing technique to screen a large number of DEGs, and exploit a large number of genes related to fatty acid anabolism.

We report the results of deep sequencing aimed at obtaining transcript coverage of Oviductus Ranae and skin obtained from *R. chensinensis* using the Illumina high-throughput sequencing platform. This technology has been widely used in various animals to obtain transcript coverage even in the absence of a reference genome. Although it has been applied to *R. chensinensis*, the purpose of this study was to analyze differences between samples based on database sequencing. According to the NR classification, more unigenes were similar to *Silurana tropicalis* and the African clawed frog because of their closer phylogenetic relationship and their abundant genomic information. In addition, the genomic information of the amphibians is not sufficiently rich. Thus, the remaining 26% of the matched genes show similarities with other species. Therefore, the unigenes in the Oviductus Ranae and skin of *R. chensinensis* should be further annotated with published gene sequences, and provide more genetic background information. The transcriptome data of *R. chensinensis* were sorted and analyzed, and 12 genes involved in the synthesis of unsaturated fatty acids were identified. Key enzyme genes involved in the synthesis of unsaturated fatty acids were also identified from the KEGG metabolic pathways. Two key enzyme genes, namely Δ6 FADS and Δ9 FADS, were enriched in the synthesis pathway of n-3 unsaturated fatty acids, while Δ5 FADS, Δ6 FADS, and Δ9 FADS were enriched in the synthesis pathway of n-6 unsaturated fatty acids. Among them, the unigene 48741 and Unigene 55182 of the noted Δ5 FADS, were annotated in the K10224 gene. Comparing Δ5 FADS with other species, Unigene55182 exhibited the highest homology and closest relationship with the sequence of the human Δ5 FADS gene. Unigene 48741 exhibited the highest homology and closest relationship with the alpine frog FADS1. We have registered the key enzyme genes screened in GeneBank to obtain the corresponding gene accession number MG879290-MG879292.

## CONCLUSION

Because of the important pharmacological effects of *R. chensinensis*, the aims of this study were to investigate the *de novo* transcriptome skin and Oviductus Ranae of *R. chensinensis* using the Illumina Hiseq 2000 platform. More importantly, on gene expression levels and identifications, functional annotations, and functional genomic studies could be explored using these transcripts. Based on the sequencing, key genes involved in biosynthesis of unsaturated fatty acids were isolated, which established a biotechnological platform for further research on *R. chensinensis*.

**SOURCE(S):** Jilin Province Key Scientific and Technological Achievements Transformation Project: 20160307004YY.

